# Bundle Analytics based Data Harmonization for Multi-Site Diffusion MRI Tractometry

**DOI:** 10.1101/2024.02.03.578764

**Authors:** Bramsh Qamar Chandio, Julio E. Villalon-Reina, Talia M. Nir, Sophia I. Thomopoulos, Yixue Feng, Sebastian Benavidez, Neda Jahanshad, Jaroslaw Harezlak, Eleftherios Garyfallidis, Paul M. Thompson

## Abstract

The neural pathways of the living human brain can be tracked using diffusion MRI-based tractometry. Along-tract statistical analysis of microstructural metrics can reveal the effects of neurological and psychiatric diseases with 3D spatial precision. To maximize statistical power to detect disease effects and factors that influence them, data from multiple sites and scanners must often be combined, yet scanning protocols and hardware may vary widely. For simple scalar metrics, data harmonization methods - such as ComBat and its variants - allow modeling of disease effects on derived brain metrics, while adjusting for effects of scanning site or protocol. Here, we extend this method to pointwise segment analyses of 3D fiber bundles by integrating ComBat into the BUndle ANalytics (BUAN) tractometry pipeline. In a study of the effects of mild cognitive impairment (MCI) and Alzheimer’s disease (AD) on 38 white matter tracts, we merge data from 7 different scanning protocols used in the Alzheimer’s Disease Neuroimaging Initiative, which vary in voxel size and angular resolution. By incorporating ComBat harmonization, we model site- and scanner-specific effects, ensuring the reliability and comparability of results by mitigating confounding variables. We also evaluate choices that arise in extending batch adjustment to tracts, such as the regions used to estimate the correction. We also compare the approach to the simpler approach of modeling the site as a random effect. To the best of our knowledge, this is one of the first applications to adapt harmonization to 3D tractometry.

## I. Introduction

In neuroimaging research, the harmonization of data acquired at different imaging sites and on different scanners is paramount to ensure the reliability, comparability, and accurate interpretation of the findings. Variations in acquisition parameters, scanner types, and imaging protocols can introduce systematic biases, making it harder to interpret and compare findings across studies. Data harmonization is crucial in addressing these challenges and enhancing the reliability and generalizability of neuroimaging research findings. Data harmonization is the process of standardizing or aligning measures from heterogeneous datasets to mitigate the impact of technical variability while maintaining the integrity of underlying biological signals. By harmonizing data across different imaging modalities, scanners, and acquisition parameters, researchers can effectively reduce unwanted sources of variability, such as scanner shifts, site-specific biases, and imaging artifacts. It enables physicians to make accurate assessments and treatment decisions.

Among several harmonization techniques, ComBat harmonization [1], [2] is a widely used method in neuroimaging, and has been used in both structural and diffusion MRI (dMRI) studies to model variations in data distributions arising from different scanning protocols, imaging sites, or scanner types. Originally designed to correct for batch effects in gene expression data, the ComBat family of tools has been widely adopted in neuroimaging. In the context of dMRI, differences in acquisition parameters (such as voxel size, angular resolution, and b-values) can influence diffusion metrics, such as fractional anisotropy (FA), mean, radial, and axial diffusivity (MD, RA, AxD) derived from diffusion tensor imaging (DTI) [3]. By standardizing diffusion metrics, researchers can more confidently conduct group-level analyses, identify subtle differences in white matter microstructure, and draw robust conclusions regarding microstructural alterations associated with various neurological or psychiatric conditions. ComBat achieves harmonization by fitting a linear model to the data, adjusting for covariates such as age and sex, and then applying a scale and shift transform to the residuals to match a reference dataset or values based on pooling all the data in a study. The scale and shift parameters (see Methods for details) are a second-order model of the variability (distributional shift) attributed to a given scanning site or protocol. When data are limited at a given site, empirical Bayes methods are used to make these batch corrections more robust.

Moreover, ComBat harmonization enables data from diverse cohorts or clinical populations to be integrated, facilitating data integration initiatives in the neuroimaging community. This fosters collaboration and accelerates scientific discovery by pooling resources and leveraging multiple existing datasets to investigate brain microstructure and its relationship to disease mechanisms.

ComBat has primarily been applied at the voxel or region of interest level to harmonize morphometric or microstructural measures when analyzing the measures for group comparisons, or as part of a multivariate statistical model. However, its application has not been well-studied for tractometry-based analysis of white matter alterations in disease. A recent paper [4] analyzed a multi-site dataset of neonates and infants with congenital heart disease and used ComBat as a post-processing step for the harmonization of along-tract data from WM bundles. However, their study focused on comparing the accuracy of manually delineated and the automated generation of each tract and along-tract. The study was focused on differences in pre- and post-operative single- and bi-ventricle surgical repair. Tractometry can combine information on brain tissue microstructural measures such as FA, MD, RD, and AxD [3] with geometric information on the shape of white matter bundles to study effects of developmental disorders such as autism, or degenerative conditions such as Alzheimer’s or Parkinson’s disease. Whole brain fiber pathways are reconstructed *in vivo* using dMRI and tractography methods, resulting in wholebrain tractograms. Particular white matter tracts of interest are then segmented from whole-brain tractograms using bundle segmentation methods [5], [6]. The microstructural measures are projected along the length of these extracted bundles, to create bundle profiles. These bundle profiles from different subjects and populations are statistically analyzed to find significant group differences at a particular location along the length of the bundles.

In this work, we incorporate the ComBat method into our established BUndle ANalytic (BUAN) tractometry pipeline [7]. We use ADNI3 (Alzheimer’s Disease Neuroimaging Initiative) [8] data from 7 unique dMRI protocols to correct for scanner acquisition protocol effects when examining the effects of mild cognitive impairment (MCI) and dementia on 38 white matter tracts. We compare bundle-wise and wholebrain approaches for feature selection for harmonization. We carefully evaluate the outcomes obtained with and without harmonization, employing various statistical approaches to examine the efficacy of harmonized BUAN tractometry in improving data consistency and detecting disease effects.

## II. Methods

Data from 730 ADNI3 participants (age: 55-95 years, 349M/381F, 214 with MCI, 69 AD, and 447 cognitively healthy controls (CN)) scanned with 7 acquisition protocols (GE36, GE54, P33, P36, S127, S31, S55) were included. Tables 1 and 2 in Fig. 2 detail demographic and acquisition protocol information. dMRI were preprocessed using the ADNI3 dMRI protocol [9], [10]. Preprocessing of raw dMRI data included denoising [11], [12], Gibbs deringing [13], [14], skull stripping [15], [16], motion and eddy current correction [16], [17], b1 bias field inhomogeneity correction [14], [18], and echo-planar imaging distortions correction. We applied robust and unbiased model-based spherical deconvolution [19] reconstruction method and a probabilistic particle filtering tracking algorithm that utilizes tissue partial volume estimation (PVE) to reconstruct [20] whole-brain tractograms. For tracking, the seed mask was created from the white matter (WM) PVE (WM PVE *>* 0.5), seed density per voxel was set to 2, and step size was set to 0.5. We extracted 38 white matter tracts from tractograms using auto-calibrated RecoBundles [7], [21] (See Fig. 2 for full names).

Here, we introduce microstructural harmonization of bundle profiles into our BUAN tractometry pipeline to correct for scanner/site effects. The harmonized BUAN pipeline steps are visualized in Fig. 1. After extracting WM bundles, BUAN creates the bundle profiles for each bundle using 4 DTI based microstructural metrics: FA, MD, RD, and AD (see 2 for full bundle names). Microstructural measures computed from different advanced modeling approaches, such as diffusion kurtosis imaging (DKI) [22], or neurite orientation dispersion and density imaging (NODDI) [23], could also be incorporated into BUAN. However, for the purpose of illustrating harmonization in tractometry, we chose these 4 well-known DTI metrics.

**Fig. 1:**
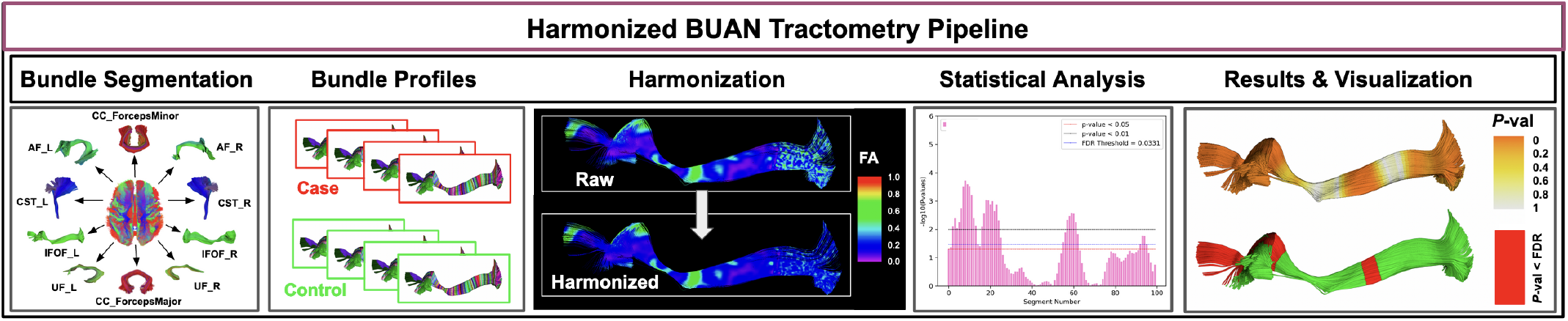
Harmonized BUAN tractometry pipeline: Bundle segmentation is performed using auto-calibrated RecoBundles, bundle profiles are created, and microstructural measures are projected onto them. Bundle profiles are harmonized using ComBat, and ComBat output is fed into Linear Mixed Models (LMMs) to discover group differences. Results are visualized both as plots and *p*-values projected onto 3D fiber bundles.

**Fig. 2:**
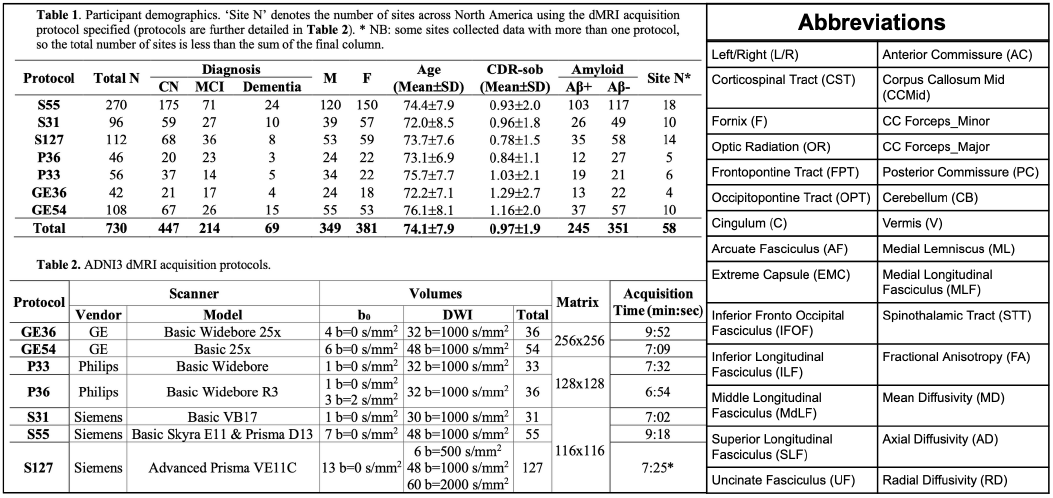
Tables 1 and 2 detail demographic and scanner protocol information for the ADNI3 data used in our experiments (data from Thomopoulos et al, 2021). The abbreviation table on the right shows the names of 38 white matter tracts and four microstructural measures analyzed in this work.

Bundle profiles are created by dividing the bundles into 100 segments using the model bundle centroids along the length of the tracts in common space. We cluster our model bundle using the QuickBundles [24] method to obtain a cluster centroid consisting of 100 points per centroid as shown in Fig. 1. We calculate Euclidean distances between every point on every streamline of the bundle and 100 points in the model bundle centroid. A segment number is assigned to each point in a bundle based on the shortest distance with the nearest model centroid point. The streamlines are not resampled to have a discrete number of points, and we do not change the distribution of points. One segment can have multiple points contributing to it from the same streamline of a subject. Since the assignment of segment numbers is performed in the common space, we establish the segment correspondence among subjects from different groups and populations. Microstructural measures such as FA are then projected onto the points of the bundles in native space. Originally, BUAN would analyze each segment for group differences using Linear Mixed Models (LMMs). To account for scanner and/or site effects, a random term is used to model them in LMMs. Here, we simplify the bundle profiles by averaging multiple points belonging to the same segment from the same subject so that each subject contributes only one value per segment. Due to limitations of the current version of ComBat, multiple values per feature cannot be handled; we decided to simplify the bundle profiles to create a central streamline with averaged values in each of 100 segments along its length. However, segments may still contain *NaN* values if no points from a streamline were assigned to them. We plan to extend ComBat for tractometry data to handle multiple values per feature in future work.

After creating bundle profiles and before running group statistics, we harmonize bundle profiles using the ComBat method to correct for scanner/site effects.

In previous studies using ROI-based ComBat harmonization, each complete ROI with a mean microstructural value associated with it is treated as one feature. Data from all ROIs are pooled together and fed into ComBat to correct for scanner/site effects, where each ROI is considered as a feature. For our tract data, we incorporate three different approaches to perform along-tract ComBat-based harmonization as described below.

### A. ComBat Tract Feature Selection

#### 1) Bundle-wise harmonization

We assume each bundle type has its own date distribution, which is considered independent of the rest of the bundles in the brain. For each tract and metric, we pool bundle profiles for a given tract across all subjects from CN, MCI, and AD groups. Pooled bundle profiles consist of 100 segments, and each segment is modeled as a feature.

#### 2) Bundle-core-wise harmonization

Bundle-core-wise harmonization is similar to Bundle-wise harmonization, except here, we first prune streamlines far from the middle of the bundle and cut *N* segments from each end of the bundle length-wise. We hypothesize that excluding the extremities of the bundles might result in more stable ComBat estimates, as the ends of the bundles tend to have varying shapes and may contain noise in the microstructural measures. This pruning and trimming process of the bundle, as visualized in Fig. 3, results in *K* segments per bundle.

**Fig. 3:**
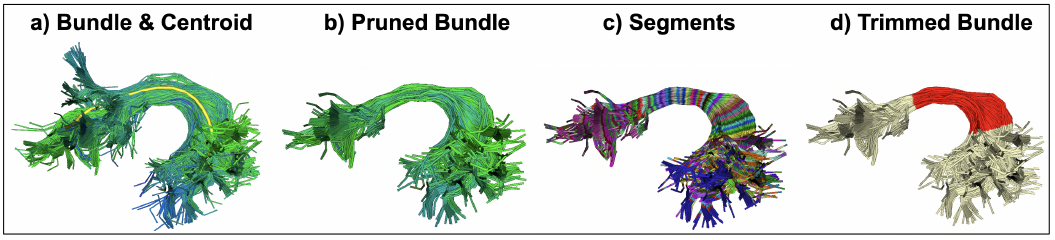
a) Model bundle centroid projected onto input bundle, (b) filtered bundle based on distance from the centroid, (c) 100 segments created along the length of the bundle, and (d) trimmed bundle to exclude bundle ends with high variation and noise. The core of the filtered and cleaned bundle is highlighted in red

After trimming down the bundle, we pool together bundle profiles for a given tract across all subjects from CN, MCI, and AD groups. Pooled bundle profiles consisting of *K* segments are modeled as features.

#### 3) Whole-brain Tractogram-wise harmonization

We pool together bundle profiles of all tracts of all the subjects coming from CN, MCI, and AD groups. The 38 pooled bundle profiles consist of 38* 100 = 3800 segments, where each segment is considered as a feature.

Data prepared using any one of the approaches described above is then fed into ComBat for scanner protocol effect correction with group, age, and sex modeled as covariates. Due to a limited number of subjects scanned at some sites in ADNI3, we chose not to model sites as nested within the protocol, as the modeling would not be robust. ComBat corrected data is then used to run a linear regression with group, age, and sex modeled as fixed effects and the response variable being a microstructural measure. Further, we use Linear Mixed Models (LMMs) where the protocol is incorporated as a random effect to find group differences between MCI and CN, AD and CN, and AD and MCI.

### B. ComBat for Tract Data Harmonization

Similar to the ComBat model [1] designed for DTI images [2], we reformulate the ComBat model for tractography-based harmonization of DTI metrics along the length of the WM tracts. Here, we use FA as a microstructural measure of interest for explaining the ComBat model.

We assume that the data come from *m* imaging protocols, containing each *n*_*i*_ scans for *i* = 1, 2, …, *m*. For segment *s* = 1, 2, …, *p*, let *y*_*ijs*_ represent the FA measure for the scan *j* at site *i*. The ComBat posits the following location and scale (L/S) adjustment model:

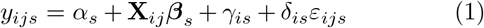

Where *α*_*s*_ is the overall FA measure for segment *s*, **X** is a design matrix for the covariates of interest (e.g. group, sex, and age), and ***β***_*s*_ is the segment-specific vector of regression coefficients corresponding to *X*. The error terms *ε*_*ijs*_ are assumed to follow a normal distribution with mean zero and variance 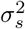. The terms *γ*_*is*_ and *δ*_*is*_ represent the additive and multiplicative scanning protocol effects of protocol *i* for segment *s*, respectively.

The procedure for the estimation of the scanner parameters *γ*_*is*_ and *δ*_*is*_ uses Empirical Bayes and is described in [1] and [2]. Briefly, it estimates an empirical statistical distribution for each of the parameters by assuming that all 100 segments along the length of the tracts share a common distribution for bundle-wise and *N* segments for bundle-core-wise harmonization. For the whole-brain tractogram-wise harmonization, 100 segments of each bundle (totaling 3800 segments) share the same distribution. Information contributed by all the segments is used to inform the statistical properties of the scanner effects and is assumed to have the parametric prior distributions: 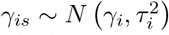 and 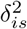 ∼ Inverse Gamma (*λ*_*i*_, *θ*_*i*_). The hyperparameters *γ*_*i*_, 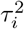, *λ*_*i*_, *θ*_*i*_ are estimated empirically from the data. The ComBat estimates 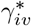 and 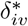 of the scanner effect parameters are computed using conditional posterior means as described in [1]. The final ComBat harmonized FA bundle profile measurements are defined as

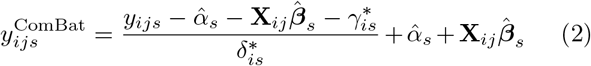

## III. Results and Discussion

Here, we compare the effect of harmonization and feature selection for harmonization on disease detection sensitivity. We ran ComBat harmonization on 38 bundles and 4 microstructural measures: FA, MD, RD, and AD. We examine the effect of harmonization on statistical analysis. We compare different bundle feature selection methods used to prepare data for ComBat. By default, harmonized BUAN uses a bundle-wise feature selection approach. For illustration purposes, most figures show results from MCI vs. CN group differences unless mentioned otherwise in the figure.

Overall, after ComBat harmonization, group differences were more sensitively detected, as compared to linear regression and LMMs with protocol modeled as a random term applied to unharmonized data. However, for some tracts, we also find comparable results when using LMMs with scanner protocol modeled as a random effect aligned with our previous work [8], [25], [26]. While harmonization enhances the significance of the group effect for most tracts, it also decreases the group difference effect for some tracts, as shown in Fig. 7, suggesting the removal of scanner-specific confounds that are correlated with the disease effect. We also visualize the results of harmonized BUAN mapping effects of MCI and AD for the left cingulum and right arcuate fasciculus (Fig. 4, bottom panel). Adding different scanner protocols in the analysis boosts disease sensitivity, and harmonization further boosts power, as shown before and after harmonization in Fig. 8f-g.

**Fig. 4:**
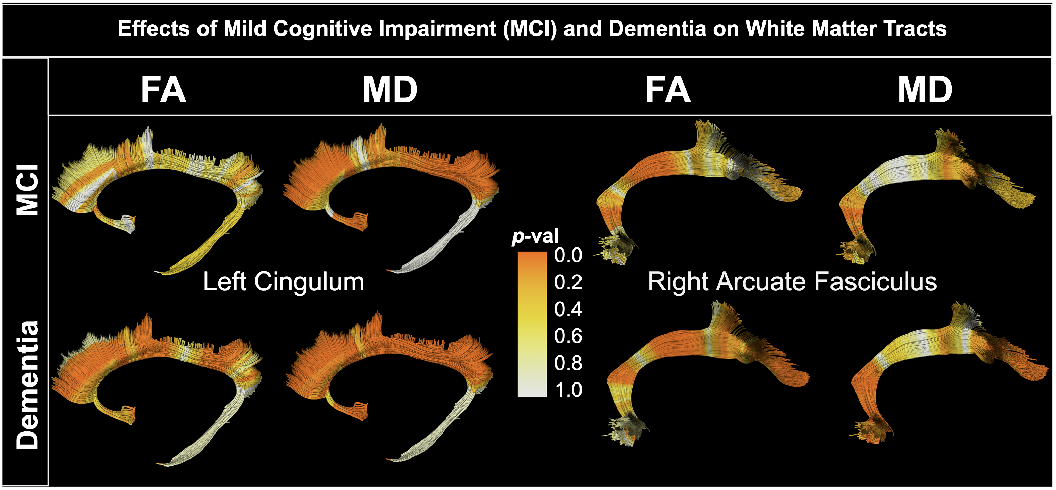
Effects of MCI and AD using the harmonized BUAN tractometry approach on the left cingulum (C_L) and right arcuate fasciculus (AF_R) bundles, for fractional anisotropy (FA) and mean diffusivity (MD) metrics.

**Fig. 5:**
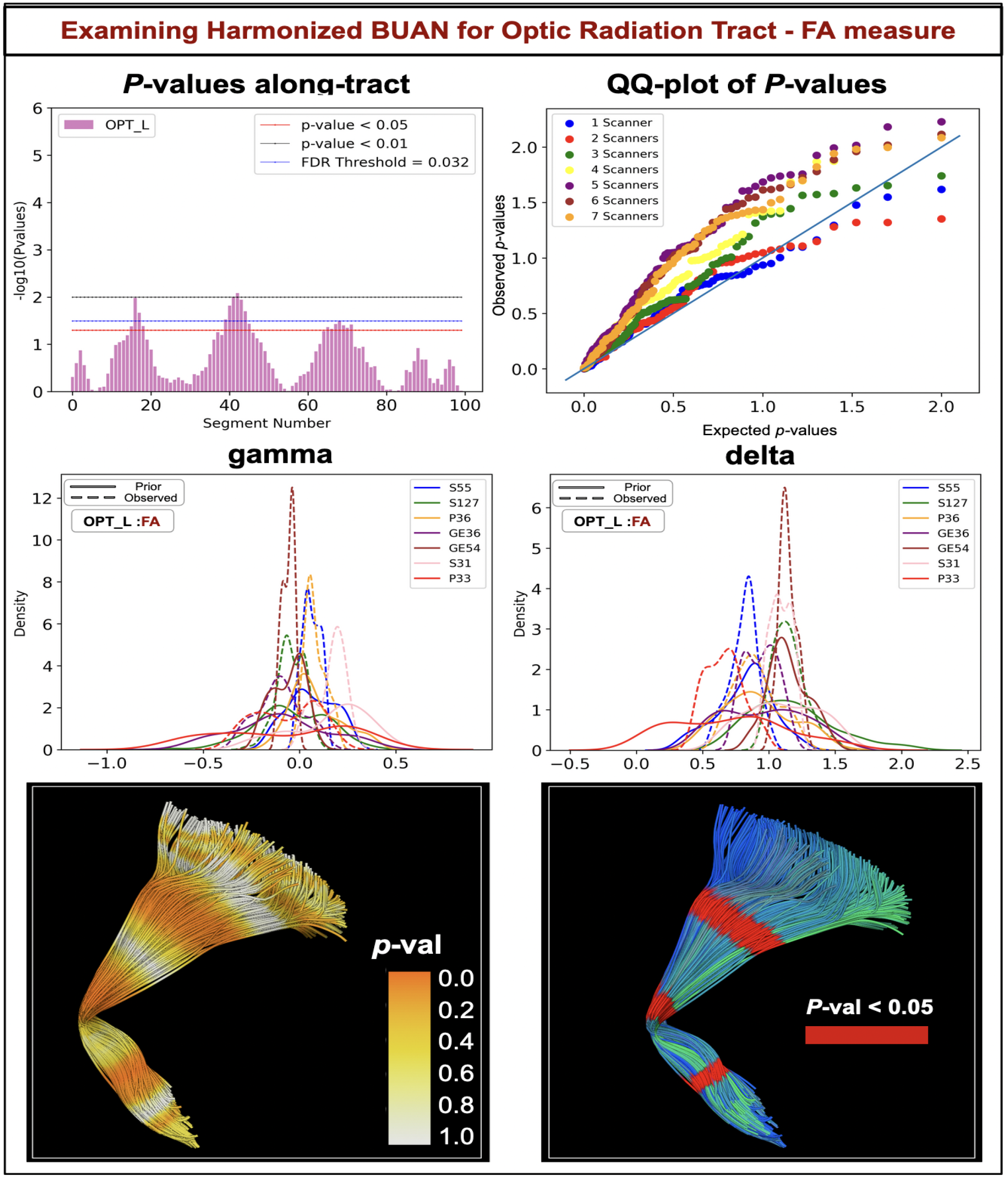
Harmonized BUAN tractometry pipeline, applied to the Optic Radiation tract in the left hemisphere of the brain (OPT_L). The top row shows a plot of the negative logarithm of *p*-values along the length of the tract and qq-plot of *p*-values, showing a boost in disease detection when data from more sites/scanners is added. The middle row shows prior and observed gamma and delta parameters in the ComBat method for the OPT_L tract. The bottom row shows *p*-values projected onto the 3D model of OPT_L tract. Significant segments are highlighted in red.

However, whole-brain tractogram-wise feature selection for harmonization reduces disease sensitivity detection as compared to bundle-wise harmonization, as shown in Fig. 7. The challenge with whole-brain tractogram-wise feature selection lies in its handling of varying data characteristics inherent when dealing with whole-brain connectivity data. The brain encompasses a diverse array of WM bundles, some of which connect to the spinal cord while others extend into gray matter regions, and the corpus callosum connects the two brain hemispheres. This inherent variability in tract composition and shape variations of tract, specifically termination points, poses a challenge for harmonization techniques that treat the entire brain tractogram as a single entity. When applying wholebrain tractogram-wise feature selection for harmonization, the method may not adequately capture the nuanced variations present across different white matter bundles. As a result, harmonization efforts may inadvertently prioritize certain bundles over others, leading to sub-optimal alignment of diffusion metrics and potentially reducing the sensitivity of disease detection.

In contrast, bundle-wise harmonization strategies offer a more targeted approach by focusing on individual white matter bundles or tracts. By treating each bundle independently during the harmonization process, these strategies may better account for the unique characteristics and variability associated with each tract. This targeted approach enhances the accuracy of harmonization and preserves the specificity of diffusion metrics within each bundle, thereby improving the sensitivity of disease detection.

In our investigation, we evaluated the efficacy of bundlecore-wise feature selection compared to the whole bundle-wise feature selection method. We expected to find an increase in disease sensitivity by excluding spurious streamlines, and especially extremities of the bundles, which tend to vary in shape. Intuitively, microstructural measures could be susceptible to confounding factors or artifacts inherent in dMRI data. Perhaps surprisingly, our findings revealed that bundlecore-wise feature selection yielded almost the same results as the whole bundle-wise approach, as shown in Fig. 6. One notable aspect of our study was the examination of how cutting segments from the ends of the tract influences sensitivity in detecting disease effects. We systematically evaluated three scenarios: utilizing the entire bundle, retaining 80% of the bundle by excising 10 segments from each end and preserving 60% of the bundle by removing 20 segments from each end of the bundle. Additionally, current work only addresses bundle-specific harmonization of microstructural measures.

**Fig. 6:**
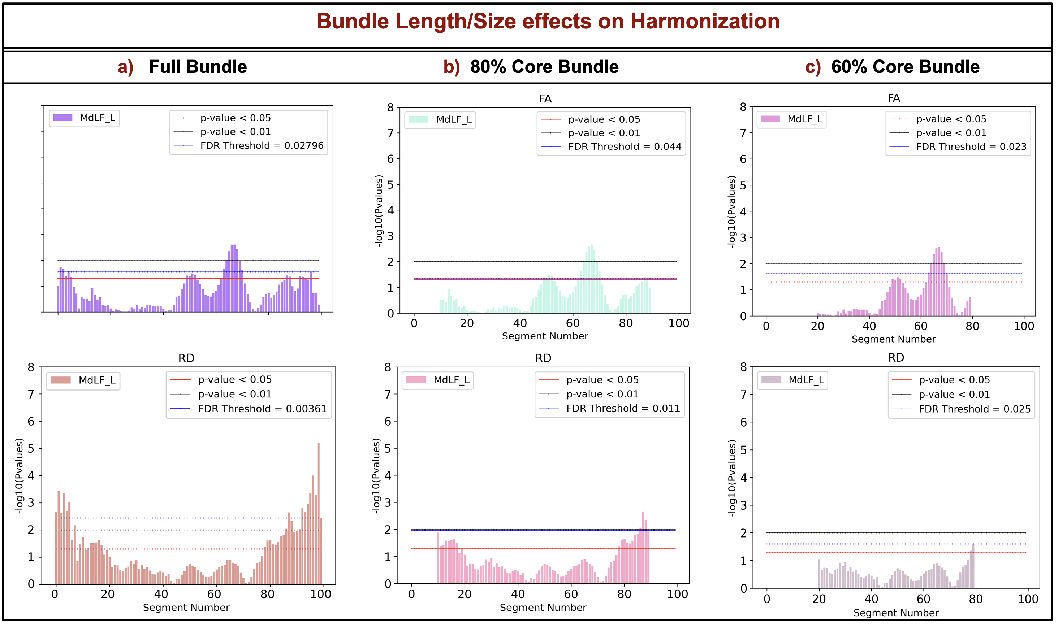
Effects of using different bundle lengths and sizes of the same bundle on the harmonization output. BUAN tractometry harmonization approach for the (MdLF_L) on fractional anisotropy (FA) and radial diffusivity (RD) metrics using different bundle coverage in statistics. We find comparable results for all 3 approaches shown in the figure, implying that keeping noisy exterminates of bundles does not cause artifacts in the statistical analysis.

**Fig. 7:**
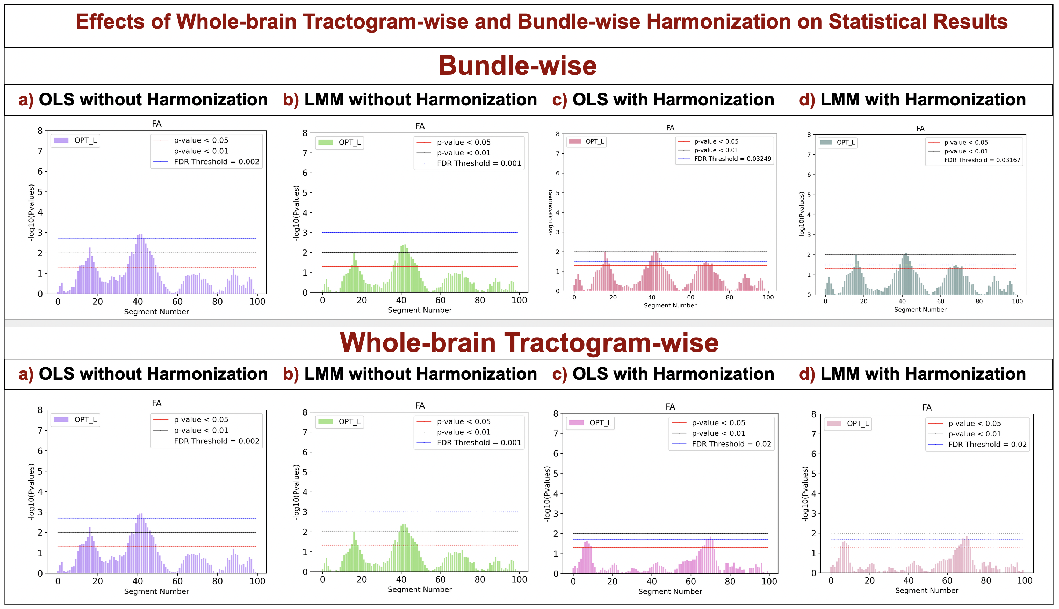
Impact of feature selection on tract microstructure harmonization: *top*, bundle-wise ComBat data harmonization and *bottom*, whole-brain tractogram-wise ComBat harmonization. Bundle-wise harmonization preserves the original data distribution and characteristics of the bundle, while whole-brain tractogram-wise harmonization may produce sub-optimal results by ignoring the tract-specific details.

**Fig. 8:**
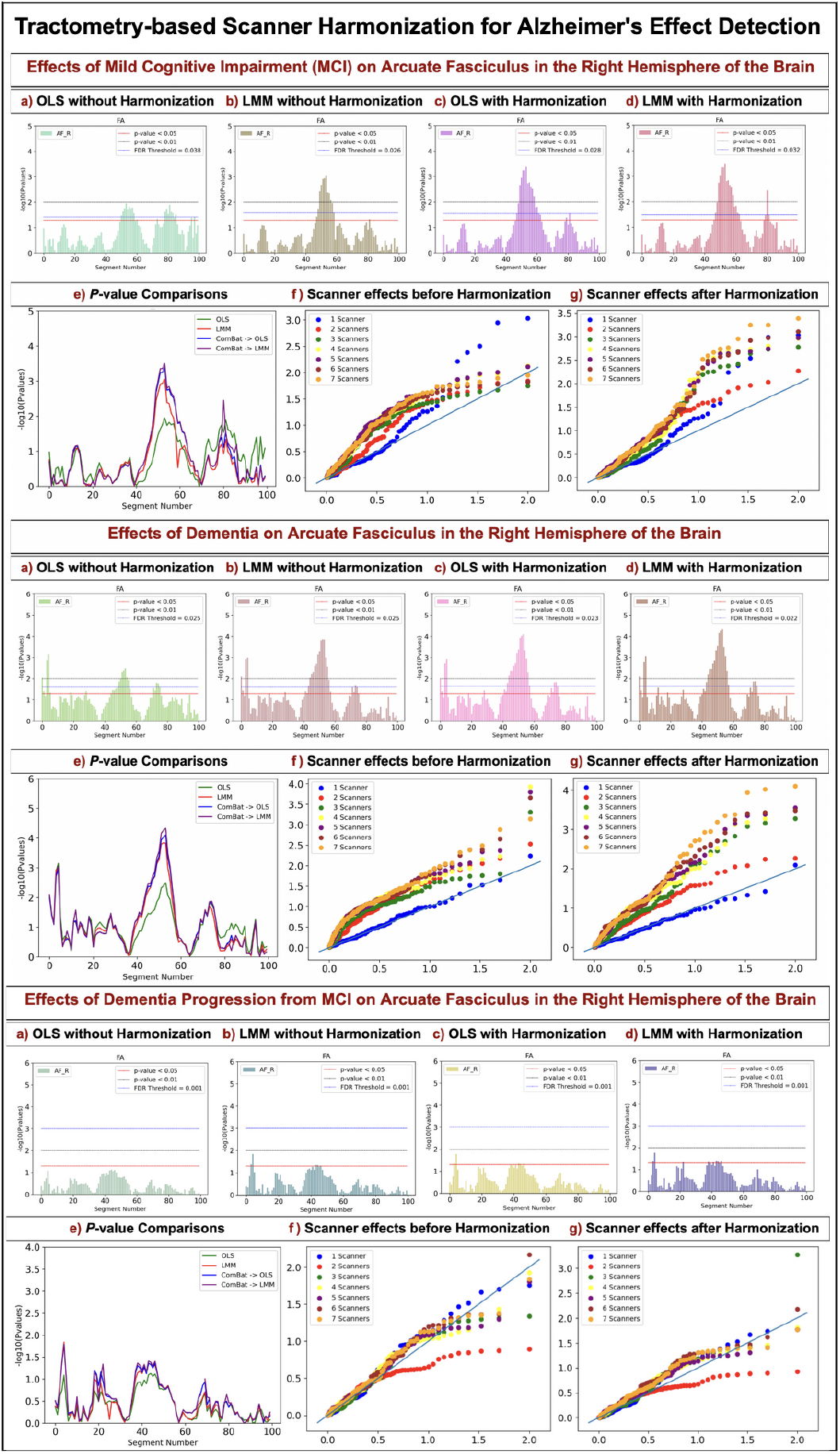
The top and bottom panels show the results for the right arcuate fasciculus (AF R) bundle for the Fractional Anisotropy (FA) measure for MCI vs. CN, AD vs. MCI, and AD vs. CN group comparisons, respectively. In all four statistical experiments, group, age, and sex were modeled as fixed effects. In both panels’ plots, the *x*-axis has segment numbers along the length of the tracts, and the *y*-axis shows the negative logarithm of the *p*-values. The FDR-corrected threshold is plotted as a blue horizontal line in the plots. a) Linear regression on unharmonized data, b) Linear Mixed Models (LMMs) with scanner protocol modeled as a random effect on unharmonized data, c) linear regression on harmonized data, and d) LMMs with scanner protocol modeled as a random effect on harmonized data. e) shows comparisons of *p*-values obtained from the four experiments. We find that linear regression and LMMs on harmonized data yield higher significance (smaller nominal *p*values), while LMMs on unharmonized data with scanner protocol modeled as random term gives comparable results. Plots f) and g) present the effects of adding data acquired from different acquisition protocols on unharmonized and harmonized data, respectively. The *x*-axis shows expected *p*-values and *y*-axis shows observed *p*-values. With each new protocol added to analyses, statistical power increases in both f) and g). While the *N* increases and this should boost power, it is possible that the protocol effect could reduce power, but this is shown to be mitigated. Harmonization in g) further helps to boost the power of analysis.

## Conclusion

Integrating ComBat into the BUAN tractometry pipeline can merge data from different sites and scanning protocols. By incorporating ComBat harmonization, we model site- and scanner-specific effects, improving the reliability and comparability of results by mitigating confounding variables. Our work has some limitations. First, ComBat is a second-order correction, whereby the mean and variance of the residuals are set to reference values, to better align data distributions. More complete alignment of data distributions is possible by either aligning higher-order moments of the residual distribution [27], or by matching the full density functions using divergence measures such as sKL or kernel-based metrics such as MMD [28]. We did not opt for these higher-order corrections, as they are prone to overfitting unless data samples are very large. Second, we are unable to fully disentangle site effects from variables that may be confounded with site. Biological measures from controls at each site are assumed to be drawn from the same underlying distribution, which may be false if sites have different inclusion criteria or co-morbidities. For this reason, a site’s data should not be aligned using ComBat unless we expect the data distribution to be similar in the controls assessed at that site. Finally, without ground truth on the geometry of the site effects, it is hard to validate that they have been completely modeled or eliminated. Empirical data on the spatial autocorrelation in the site correction fields may help determine the optimal functional form for the correction fields in the future. To the best of our knowledge, this is one of the first studies of applying harmonization to tractometry analysis. Future research will evaluate advanced ComBat variants (such as ComBat-GAM and CovBat) and advanced deep-learning approaches for distributional alignments, such as variational autoencoders (VAEs) and normalizing flows. Additionally, we aim to integrate streamline-based nonlinear bundle registration [29] to address bundle shape variability due to scanner and site differences.

## Acknowledgment

This research was supported by the NIH (National Institutes of Health) under the AI4AD project grant U01 AG068057, grant numbers P41 EB015922, and RF1 AG057892. We would like to acknowledge the National Institute of Biomedical Imaging and Bioengineering under award numbers R01 EB027585 and R01 EB017230.

